# The membrane associated accessory protein is an adeno-associated viral egress factor

**DOI:** 10.1101/2021.06.17.448857

**Authors:** Zachary C. Elmore, L. Patrick Havlik, Daniel K. Oh, Heather A. Vincent, Aravind Asokan

## Abstract

Adeno-associated viruses (AAV) rely on helper viruses to transition from latency to lytic infection. Some recombinant AAV serotypes are secreted in a pre-lytic manner as extracellular vesicle (EV)-associated particles, although mechanisms underlying such are unknown. Here, we discover that the membrane-associated accessory protein (MAAP), expressed from a (+1) frameshifted open reading frame (ORF) in the AAV capsid (*cap*) gene, is a novel viral egress factor. MAAP contains a highly conserved, cationic amphipathic domain critical for AAV secretion. Wild type or recombinant AAV with a mutated MAAP start site (MAAPΔ) show markedly attenuated secretion and correspondingly, increased intracellular retention. Trans-complementation with recombinant MAAP restored extracellular secretion of multiple AAV/MAAPΔ serotypes. MAAP is sorted into recycling Rab11+ vesicles and strongly associates with EV markers upon fractionation. In addition to characterizing a novel viral egress factor, these studies highlight a prospective engineering platform to modulate secretion of AAV vectors or other EV-associated cargo.

## Introduction

Adeno-associated virus (AAV) is a non-enveloped, single-stranded DNA virus belonging to the Dependoparvovirus genus within the Parvoviridae family^1^. Upon co-infection with a helper such as Adenovirus, Herpesvirus or Papillomavirus, AAV undergoes a transition from latent to lytic life cycle, hijacking the host cell machinery^1^. The AAV capsid consists of 60 capsid monomers of VP1, VP2, and VP3 at a ratio of 1:1:10 that packages a 4.7-kb single-stranded genome^2^. The AAV genome encodes replication (*rep*), capsid (*cap*), and assembly-activating protein (AAP) open reading frames (ORFs) flanked by inverted terminal repeats (ITRs), which are the sole requirements for genome packaging^3–5^. As such, the majority of the genome can be replaced by exogenous DNA sequences and packaged inside the AAV capsid to create a recombinant vector both *in vitro* and *in vivo*^6,7^. Some recombinant AAV serotypes appear to be secreted into cell culture media prior to lysis, albeit with variable efficiency^8–10^. Exactly how AAV exits the cell upon transitioning into this phase of replication remains unclear. Unlike other autonomous parvoviruses that undergo a lytic cycle, wild type AAV does not induce marked cytopathic effects (CPE) and therefore, cellular egress of virions is thought to be primarily driven by overexpression of Adenoviral or Herpesvirus proteins^11,12^.

Multiple studies have demonstrated that a significant fraction of recombinant AAV associate with extracellular vesicles (EVs) and are released into the supernatant fraction of the cell culture media^13^. It is well known that cells shed a variety of membrane-bound vesicles varying in size from 20 nm to 1 μm in diameter, which have been termed exosomes, microparticles or microvesicles (collectively referred to here as extracellular vesicles or EVs). Such EVs can package different macromolecules including proteins, nucleic acids and viruses, thereby making them an attractive therapeutic platform^14^. Despite being non-enveloped viruses, recombinant AAV capsids associated with EVs can enable efficient gene transfer to the retina, the nervous system, the inner ear^15–17^ and appear shielded from anti-AAV neutralizing antibodies^17^. However, the exact mechanism by which AAV associates with EVs remains to be determined.

Recent work has revealed a novel +1 frameshifted open reading frame (ORF) in the VP1 region of the AAV *cap* gene that mediates expression of the membrane-associated accessory protein (MAAP) (**Fig. 1A**), which was postulated to limit AAV production through competitive exclusion^18^. Here, we assign a novel function to MAAP in promoting EV mediated AAV egress from host cells. Specifically, we dissect the secondary structure elements in MAAP that contribute to this secretory function. Further, we demonstrate that MAAP transcomplementation can not only rescue but significantly alter the kinetics of viral secretion of multiple AAV serotypes. Further, independent of AAV infection, MAAP alone appears to possess a unique ability to induce hyper-secretion of homogenous EV fractions. Our findings highlight a viral egress mechanism evolved by AAV as well as highlight the potential to exploit this novel protein for loading therapeutic cargo including AAV vectors into EVs and AAV vector production in general.

**Figure 1.**
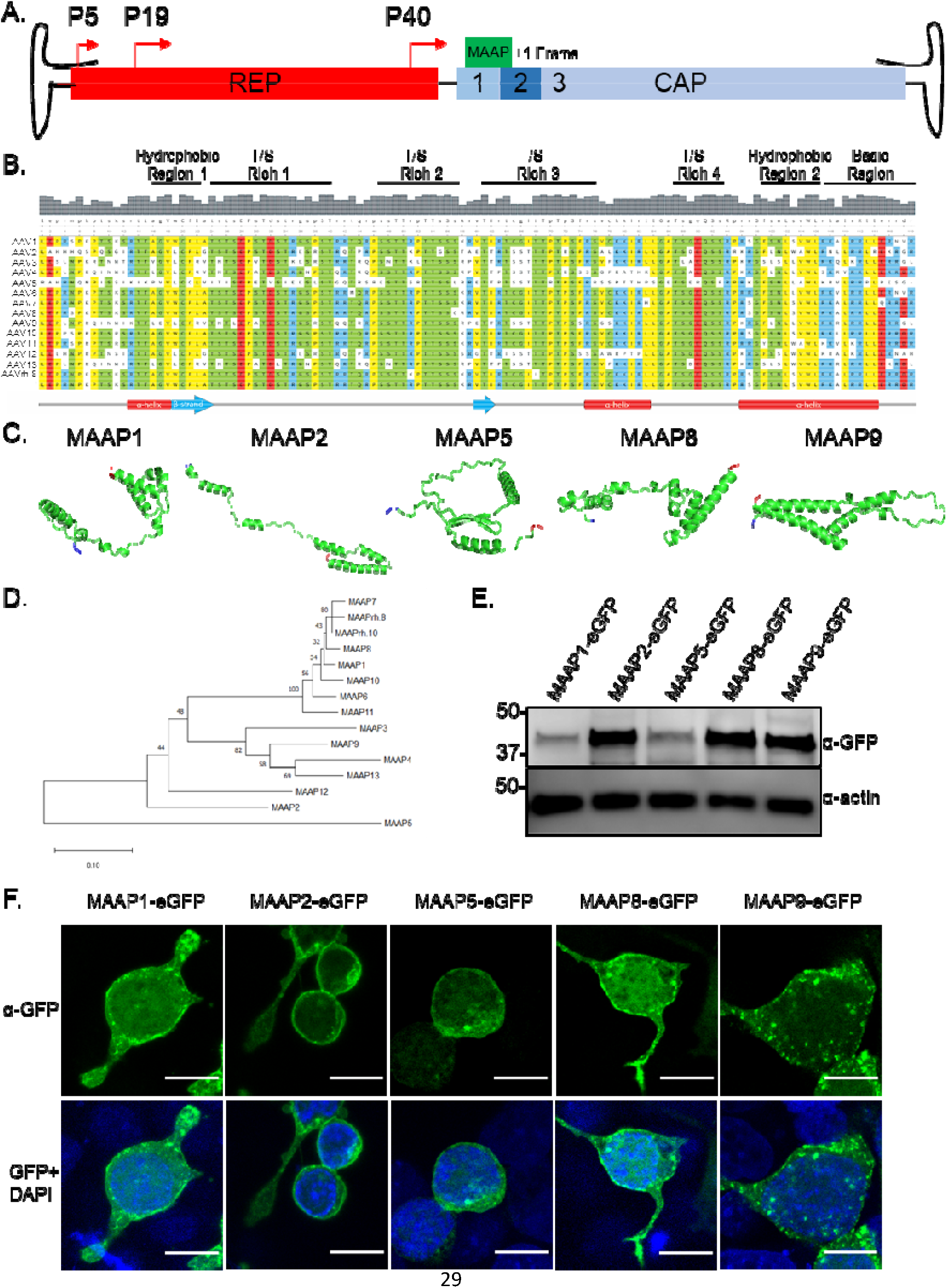
MAAP is a unique virally encoded protein with an amphipathic, cationic membrane anchoring domain. **(A)** WT AAV genome showing *Rep* and *Cap* genes with MAAP encoded in a +1 open reading frame in the VP1 region. **(B)** Sequence alignment of the MAAPs from AAV serotypes 1 to 13 along with AAVrh.8 and AAVrh.10. An annotated multiple-sequence alignment of the AAP sequences of 15 AAV serotypes is shown. Coloring reflects the physicochemical properties of the residues (Yellow=hydrophobic, Green=polar, Blue=basic, Red=acidic). Regions of interest are annotated above the alignment. Predicted secondary structural (SS) elements (strand, helix) for the sequence and amino acid numbering are displayed below the alignments. **(C)** Structural models of MAAP1, MAAP2, MAAP5, MAAP8 and MAAP9 generated using Phyre^2^ protein modeling software. Residues highlighted in blue indicate N-terminus and residues in red indicate C-terminus. **(D)** Neighbor-joining phylogeny of MAAP amino acid sequences from AAV serotypes 1 to 13, MAAPrh.8 and MAAPrh.10. MAAP amino acid sequences were aligned with ClustalW, the phylogeny was generated using a neighborjoining algorithm, and a Poisson correction was used to calculate amino acid distances, represented as units of the number of amino acid substitutions per site. The tree is drawn to scale, with branch lengths in the same units as those of the evolutionary distances used to infer the tree. Bootstrap values were calculated with 1,000 replicates, and the percentage of replicate trees in which the associated taxa clustered together are shown next to the branches. **(E)** Anti-GFP immunoblot of whole-cell extracts prepared from HEK293 cells expressing indicated GFP tagged constructs. Anti-actin immunoblot served as loading control. **(F)** Confocal images of HEK293 cells overexpressing eGFP tagged MAAP constructs. Scale bar=10μM.

## Results

### MAAP is a conserved AAV *cap* encoded protein with a predicted C-terminal membrane anchoring domain

Confirming that MAAP is a novel viral-encoded protein of unknown function, a pBLAST search of multiple AAV *cap* gene derived MAAP sequences on the National Center for Biotechnology Information (NCBI) website did not return any proteins with significant homology^18^. Amino acid sequence alignment of MAAPs derived from different AAV serotypes revealed conserved N- and C-terminal regions containing hydrophobic and basic amino acid residues interconnected by a threonine/serine (T/S) rich region (**Fig. 1B**). MAAP 3D structures generated using TrRosetta deep learning based modeling^19^ predicted the following, (i) a conserved N-terminal hydrophobic motif with both alpha helical and beta strand secondary structure elements; (ii) four T/S rich sequence clusters spanning 7-17 residues in length with last two being separated by a smaller alpha helical interspersed with basic residues and (iii) a C-terminal domain defined by another hydrophobic alpha helical motif merging into a cluster of arginine/lysine (R/K) residues (**Fig. 1B, C**). Importantly, secondary structure software analysis strongly predicts that the C-terminal domain constitutes a putative membrane binding, cationic amphipathic peptide (residues 96-114). It is noteworthy to mention that the secondary structure of MAAP is strikingly similar to the assembly activating protein (AAP), which is similarly encoded downstream from a (+1) frameshifted ORF in the *cap* gene. When combined with phylogenetic analysis using the neighbor-joining tree method, we observed that MAAPs from AAV serotypes 1,6,8,10,11 were tightly clustered, while other sequences, in particular, MAAP2, 5 and 9 showed significant divergence from other serotypes (**Fig. 1D**). We then transfected plasmids encoding recombinant MAAPs derived from the VP1 sequences of AAV serotypes 1, 2, 5, 8, and 9 and fused to a C-terminal green fluorescent protein (GFP) *in vitro* to assess their expression (**Fig. 1E**) and cellular localization. Fluorescence micrographs confirmed the propensity of MAAP to associate with cell surface membranes as well as subcellular organelles as observed by the punctate patterns throughout the cell (**Fig. 1F**). Taken together, these data confirm that MAAP is a novel AAV protein predicted to contain cationic amphipathic C-terminal domain potentially critical for membrane anchoring.

### MAAP is essential for extracellular secretion of wild type and recombinant AAV particles

We then sought to determine whether MAAP played a role in the synthesis of (i) (pseudo)wild type AAV serotype 8 (i.e., wtAAV8 packaging AAV2 *rep* and AAV8 *cap* flanked by AAV2 inverted terminal repeats [ITRs])) (**Fig. 2A**) and (ii) recombinant AAV8 (i.e., rAAV8 packaging a chicken beta-actin promoter driven luciferase transgene flanked by AAV2 ITRs) (**Fig. 2D**). To ablate MAAP expression, we mutated the CTG start codon in the MAAP alternative open reading frame (ORF) without affecting the VP1 ORF in both wtAAV8 and rAAV8 plasmids. Culture media and cell pellets were harvested following co-transfection with an Adenovirus helper plasmid (and an additional ITR flanked luciferase encoding transgene cassette in case of rAAV) on days 3 and 5. Strikingly, quantitative PCR of viral genomes revealed a significantly higher (~1 log) amount of extracellular wtAAV8 particles in contrast to MAAPΔ particles recovered from media on day 3 (**Fig. 2B**). Further, we observed delayed secretion in case of MAAPΔ particles, which were equally apportioned between extracellular and cell lysate fractions on day 5. Although statistically significant, overall viral titers on day 5 were only minimally altered. Of the total virus produced, nearly 70% of wtAAV8 particles were secreted by day 3, while MAAPΔ particles recovered in the extracellular fraction comprised <10% of total (**Fig. 2C**).

**Figure 2.**
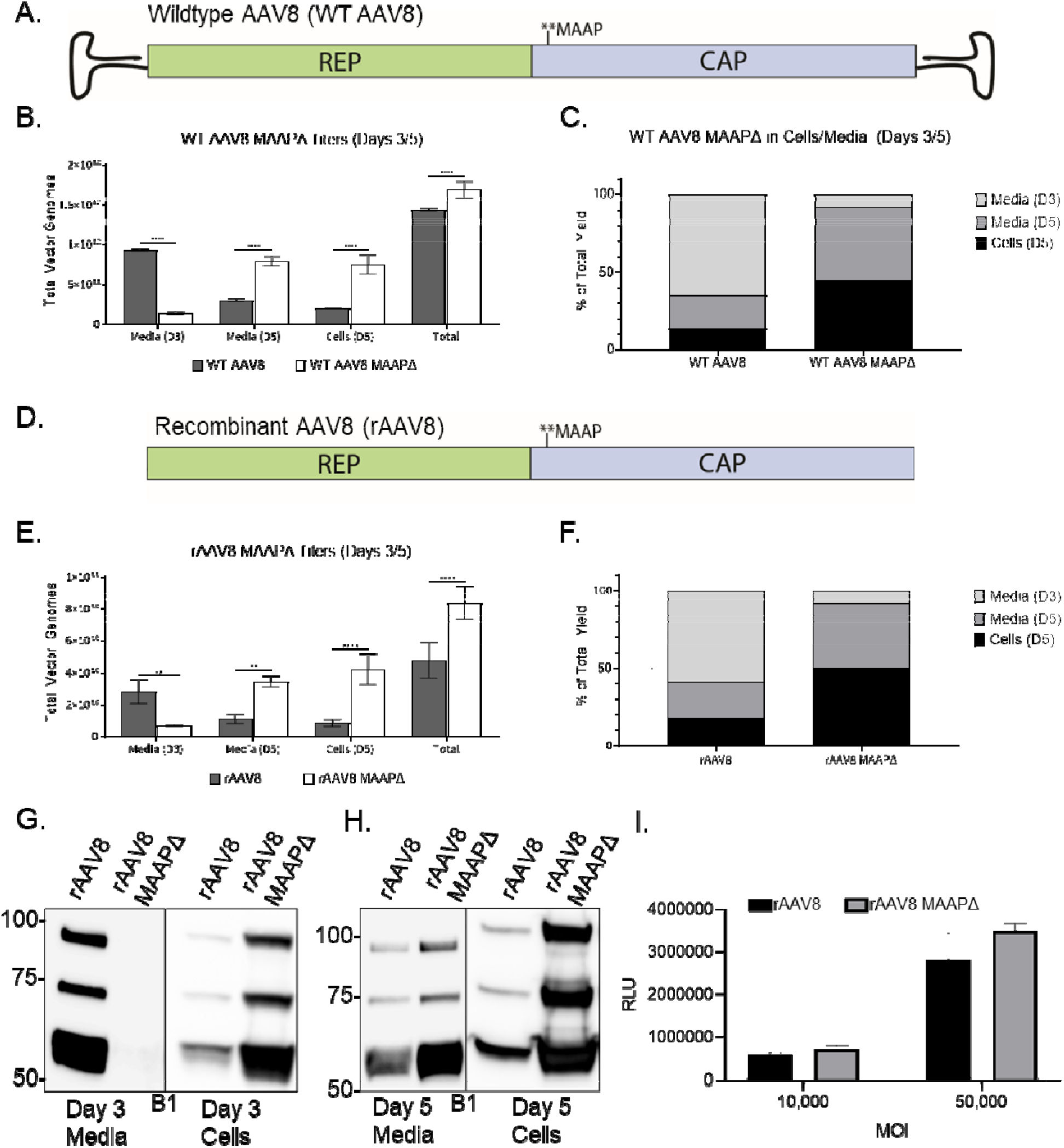
Ablation of MAAP expression results in a significant delay in the extracellular secretion of wildtype and recombinant AAV8 particles. **(A)** Schematic of WT AAV8 MAAPΔ mutant. **(B)** AAV8 ssCBA-Luc vectors produced with WT Cap or MAAPΔ Cap. Total vector genomes collected from the cells and media and the proportion of virus found in each media harvest or associated with the cells **(C)** are shown. **(D)** Schematic of rAAV8 MAAPΔ mutant. **(E)** AAV8 ssCBA-Luc vectors produced with recombinant Cap or MAAPΔ Cap. Total vector genomes collected from the cells and media and the proportion of virus found in each media harvest or associated with the cells **(F)** are shown. Recombinant AAV8 and AAV8 MAAPΔ viruses were analyzed from the media and pellet of HEK293 producing cells at days 3 **(G)** and 5 **(H)** post transfection. Capsid proteins were analyzed by SDS-PAGE under reducing conditions and probed with a capsid (B1) specific antibody. **(I)** Luciferase assay analyzing transduction of HEK293 cells by AAV8 and AAV8 MAAPΔ mutant virus at MOIs of 10,000 and 50,000 vg/cell. Each bar is a representation of three experiments that are biological replicates. Error bars indicate standard deviation from the mean. Significance was determined by two-way ANOVA, with Sidak’s post-test. \**p*<0.05, \*\**p*<0.01, \*\*\**p*<0.001, \*\*\*\**p*<0.0001.

A similar trend was observed in case of rAAV8 particles, with a 4-5 fold higher recovery from media over cell lysate and delayed secretion in case of MAAPΔ particles (**Fig. 2E**). Of the total virus produced, ~60% of rAAV8 particles were secreted by day 3 in contrast to <10% of MAAPΔ particles (**Fig. 2F**). Further evaluation of AAV capsid proteins – VP1,2 and 3 by western blot confirmed these results with undetectable to relatively lower levels in the extracellular fraction on days 3 and 5, respectively and correspondingly high(er) cellular retention in case of MAAPΔ particles (**Fig. 2G,H**). No differences were observed when comparing the transduction efficiency of rAAV8 and MAAPΔ particles *in vitro* (**Fig. 2I**). Furthermore, MAAPΔ recombinant virus showed similar VP1, VP2, and VP3 expression ratios and overall virus morphology compared to rAAV8 (**Supplementary Fig. 1**). Taken together, these results suggest that encoding MAAP from the alternative ORF in VP1 is essential for efficient cellular egress of AAV particles. Another interesting observation is the relatively low recovery of recombinant AAV serotype 9 (rAAV9) (~15%) and MAAPΔ particles (~5%) from media on day 3 (**Supplementary Fig. 2**). In contrast to rAAV8, rAAV9 particles appear to display delayed secretion as reported previously^20^, with only a modest difference in cellular egress efficiency compared to MAAPΔ particles. Thus, MAAP8 has a higher propensity to promote viral egress when compared to MAAP9. When taken together with the differences in sequence homology between AAV8 and AAV9, it is tempting to speculate that the cellular egress efficiency of different AAV capsids could be determined by their cognate MAAPs.

### Helical and cationic domains are indispensable for MAAP expression and supporting AAV secretion

We next sought to determine the regions and/or domains of MAAP necessary to promote AAV viral egress. We first inserted a 3X-FLAG tag onto the C-terminus of MAAP8 under the control of its endogenous promoter and verified its expression and function (**Supplementary Fig. 3**). At the primary structure level, MAAP can be separated into multiple regions including N-terminus, Linker, C-terminus, hydrophobic regions 1 and 2, threonine/serine-rich regions 1-4, (T/S), and the basic region. Taking a structure function approach, we generated multiple MAAP8 deletion mutants to systemically dissect the regions of MAAP critical for AAV viral egress (**Fig 3A**). We discovered that deletions of the N or C-terminus severely diminished MAAP expression (**Fig. 3B,C**). MAAP deletions involving alpha helical domains or amphipathic domains rich in basic residues demonstrated the most deleterious effects on protein expression at steady state. Furthermore, of the MAAP8 deletion mutants that expressed, only T/S-rich region 2 and T/S-rich region 4 (MAAP8 Δ3 and MAAP8 Δ5 respectively) were found to be dispensable for MAAP function as measured through AAV secretion into the media (**Fig. 3D-G**). Taken together, these data identify critical secondary structure elements in MAAP that are likely essential for expression and function.

**Figure 3.**
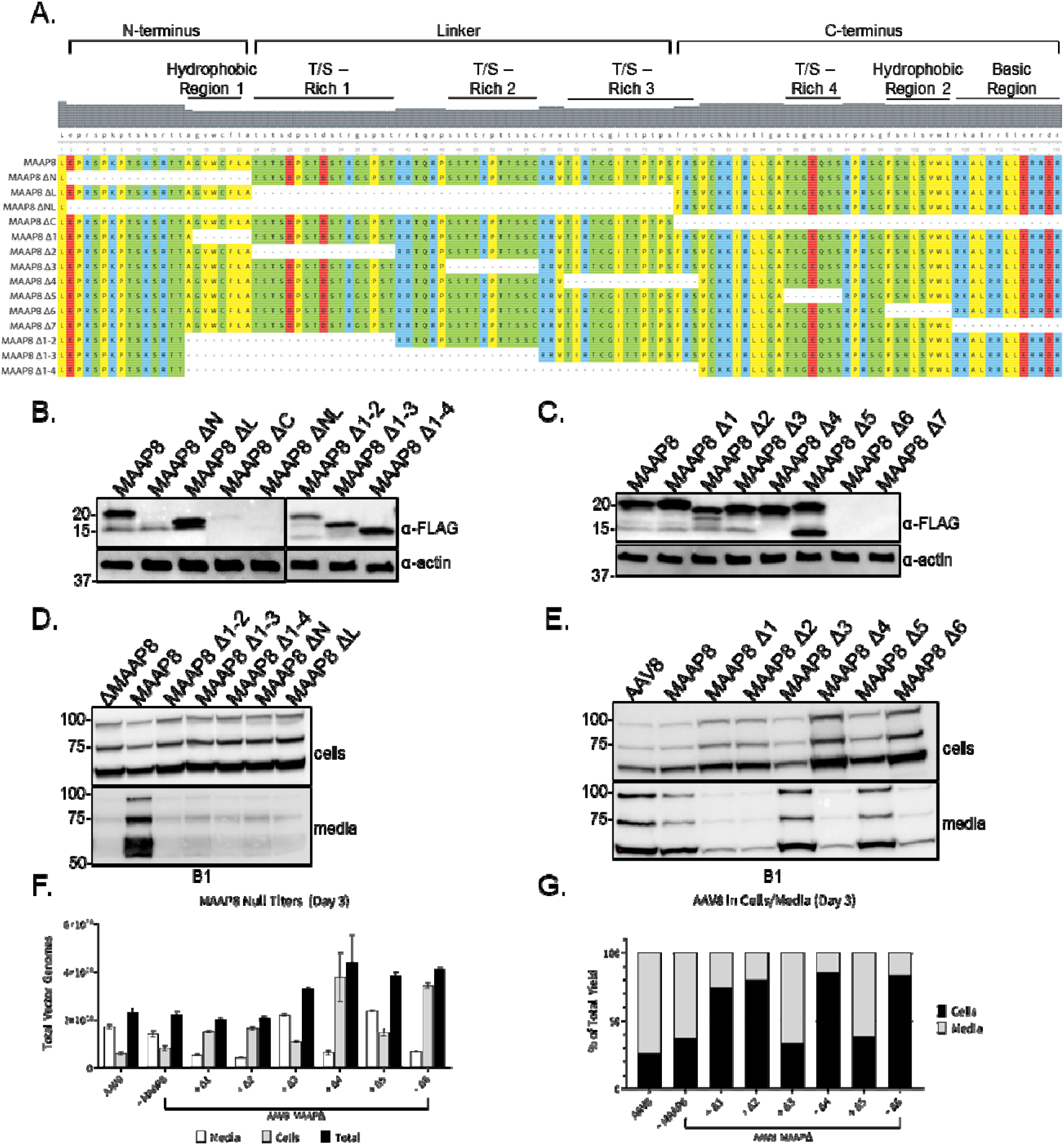
Structure-function analysis of MAAP8 reveals regions critical for expression and AAV secretion. **(A)** Schematic of different MAAP mutants. All MAAP mutants have a 3X-FLAG tag at the C terminus. **(B, C)** Anti-FLAG immunoblot of whole-cell extracts prepared from HEK293 cells expressing indicated MAAP8-3X-FLAG tagged constructs. Anti-actin immunoblot served as loading control. **(D, E)** Recombinant MAAP8Δ vectors complemented *in trans* with various truncated MAAP8-3X-FLAG plasmids were analyzed from the media and pellet of HEK293 producing cells at day 3 post transfection. Capsid proteins were analyzed by SDS-PAGE under reducing conditions and probed with a capsid (B1) specific antibody. Total vector genomes **(F)** and the proportion of vector found in the media and cells **(G)** 3 days posttransfection. Each bar is a representation of three experiments that are biological replicates. Error bars indicate standard deviation from the mean.

### MAAP transcomplementation promotes extracellular secretion of diverse AAV serotypes

To determine whether MAAP expression regulates the secretion of other AAV serotypes, we mutated the CTG start codon in the MAAP ORF for rAAV serotypes 1, 2, 8, and 9, and compared viral titers in extracellular and cellular fractions at day 3 as described earlier. In parallel, we also evaluated whether MAAP transcomplementation could rescue the extracellular secretion of MAAPΔ rAAV particles. To achieve the latter, we expressed MAAP alone from the AAV helper plasmid containing *rep* and *cap* genes by mutating the start codons in the VP1,2,3 as well as AAP ORFs. Strikingly, viral titers associated with the cellular fraction were markedly increased for rAAV1, rAAV8 and rAAV9 (~4 to 7 fold), but not rAAV2 (**Fig. 4A, C, E, G**). In addition, overall recovered titers were increased moderately for the same serotypes (up to 2 fold). In corollary, we observed a striking impact of ablating or supplementing MAAP expression on extracellular vs cell-associated fractions of different AAV serotypes. Specifically, in case of rAAV1, this percentage was reversed from 60:40 to 20:80 upon MAAP ablation and restored to normal upon MAAP expression (**Fig. 4B**). A similar trend was observed for rAAV8 (~80:20 to 20:80 followed by restoration to normal upon supplementation) (**Fig. 4F**). Both rAAV2 and rAAV9 showed decreased secretion in general (~35:65) compared to rAAV1/8 with MAAP ablation further reducing extracellular viral titers to 15% and 20%, respectively (**Fig. 4D, H**). Another important observation is that MAAP8 *trans*-complementation not only fully rescued the extracellular secretion of rAAV1, rAAV2, and rAAV8 particles, but also doubled the recovery of rAAV9 MAAPΔ particles from media compared to rAAV9 particles (from 40% to 80%) (**Fig. 4H**). These results confirm the critical role played by MAAP in enabling extracellular secretion of AAV particles in a serotype-independent manner, albeit with different efficiencies. Further, our results demonstrate that *trans*-complementation of MAAP derived from AAV8 can not only rescue secretion of different AAV serotypes, but also potentially enhance the kinetics of secretion.

**Figure 4.**
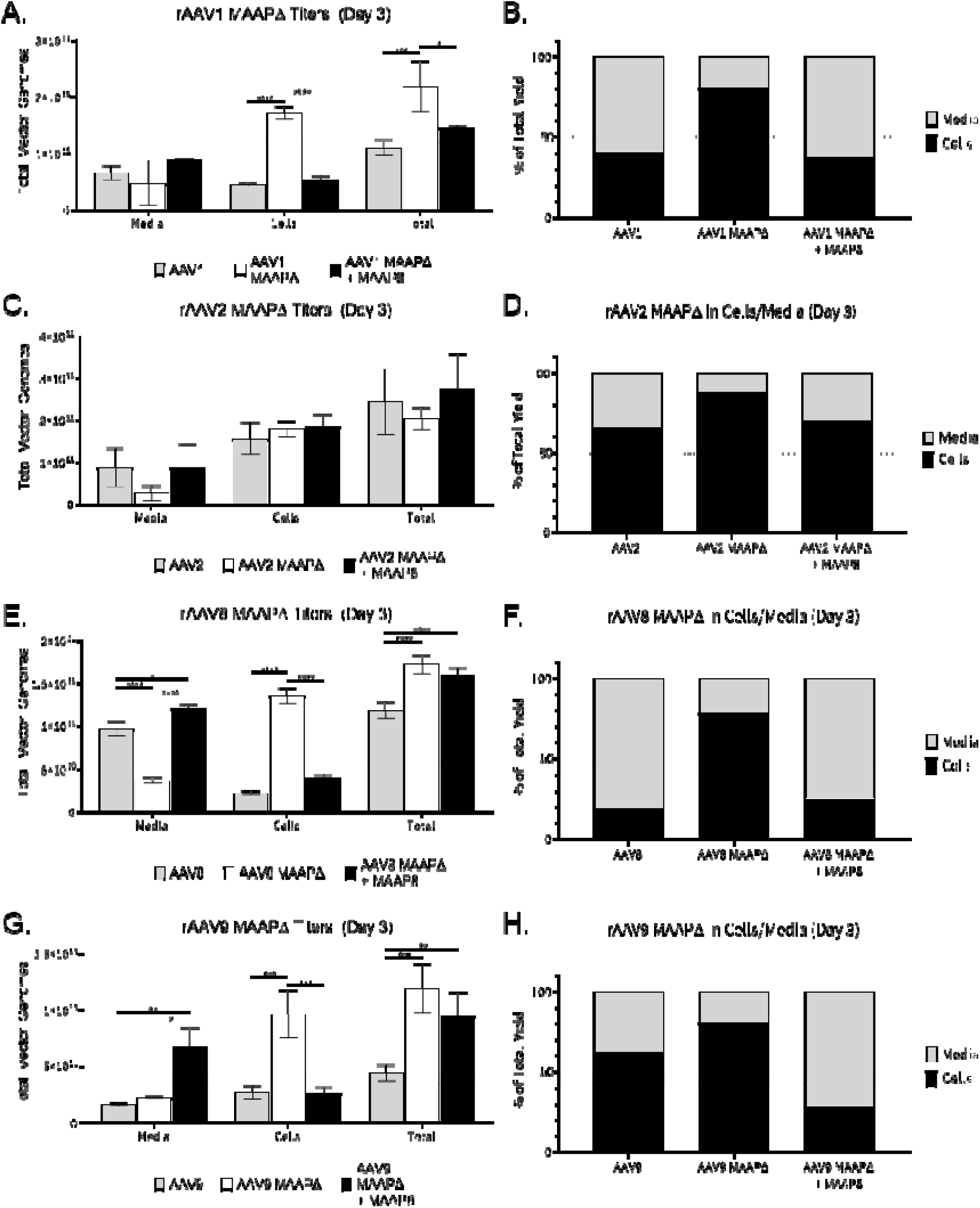
Trans-complementation of MAAP8 rescues secretion defect across multiple MAAPΔ AAV serotypes. scCBh-GFP vectors produced with WT Cap or MAAPΔ Cap for AAV1 **(A, B)**, AAV2 **(C, D)**, AAV8 **(E, F)**, and AAV9 **(G, H)** complemented *in trans* with a VP/AAP-null AAV8 plasmid to replicate endogenous levels of MAAP expression. Total vector genomes **(A, C, E, G)** and the proportion of vector found in the media and cells **(B, D, F, H)** 3 days post-transfection. Each bar is a representation of three experiments that are biological replicates. Error bars indicate standard deviation from the mean. Significance was determined by two-way ANOVA, with Tukey’s post-test. \**p*<0.05, \*\**p*<0.01, \*\*\**p*<0.001, \*\*\*\**p*<0.0001.

### MAAP enables AAV egress by hijacking vesicular secretion pathways

To obtain further mechanistic insight into how MAAP enables AAV cellular egress, we first carried out quantitative confocal fluorescence microscopy of cells overexpressing hemagglutinin (HA) tagged MAAP8, which was confirmed separately by western blot analysis (**Supplementary Fig. 4**). MAAP8-HA colocalized significantly more with the exosomal biogenesis pathway marker Rab11^21–23^, than the late endo/lysosomal pathway marker Rab7^24^ (**Fig. 5A,B**). Interestingly, MAAP9, which was associated earlier with slower secretion kinetics showed a similar co-localization to both Rab7 and Rab11 subcellular compartments (**Supplementary Fig. 5**). Overall, these results underscore the finding that MAAP is a key viral factor that mediates cellular egress of AAV particles through EV secretion pathways, an observation reported by others^13,15–17,25,26^. Further, our results also highlight potential differences in the structural attributes of different MAAPs that may explain the ability to exploit distinct secretory mechanisms that enables cellular egress of different AAV serotypes. Although the exact components of the vesicular secretion pathway exploited in MAAP-aided AAV egress still need to be elucidated, we carried out biochemical assays to explore potential MAAP-AAV capsid interactions. Notably, we did not find any evidence of direct interaction between MAAP and AAV capsid proteins or MAAP and AAP as determined by immunoprecipitation analysis (**Supplementary Fig. 6A,B**). However, it is important to note that MAAP8 fused to the BirA biotin ligase BioID2^27^ was able to successfully biotinylate the AAV capsid proteins *in vitro* indicating that AAV and MAAP do share a proximal interaction within the cell (**Supplementary Fig. 6C-F**).

**Figure 5.**
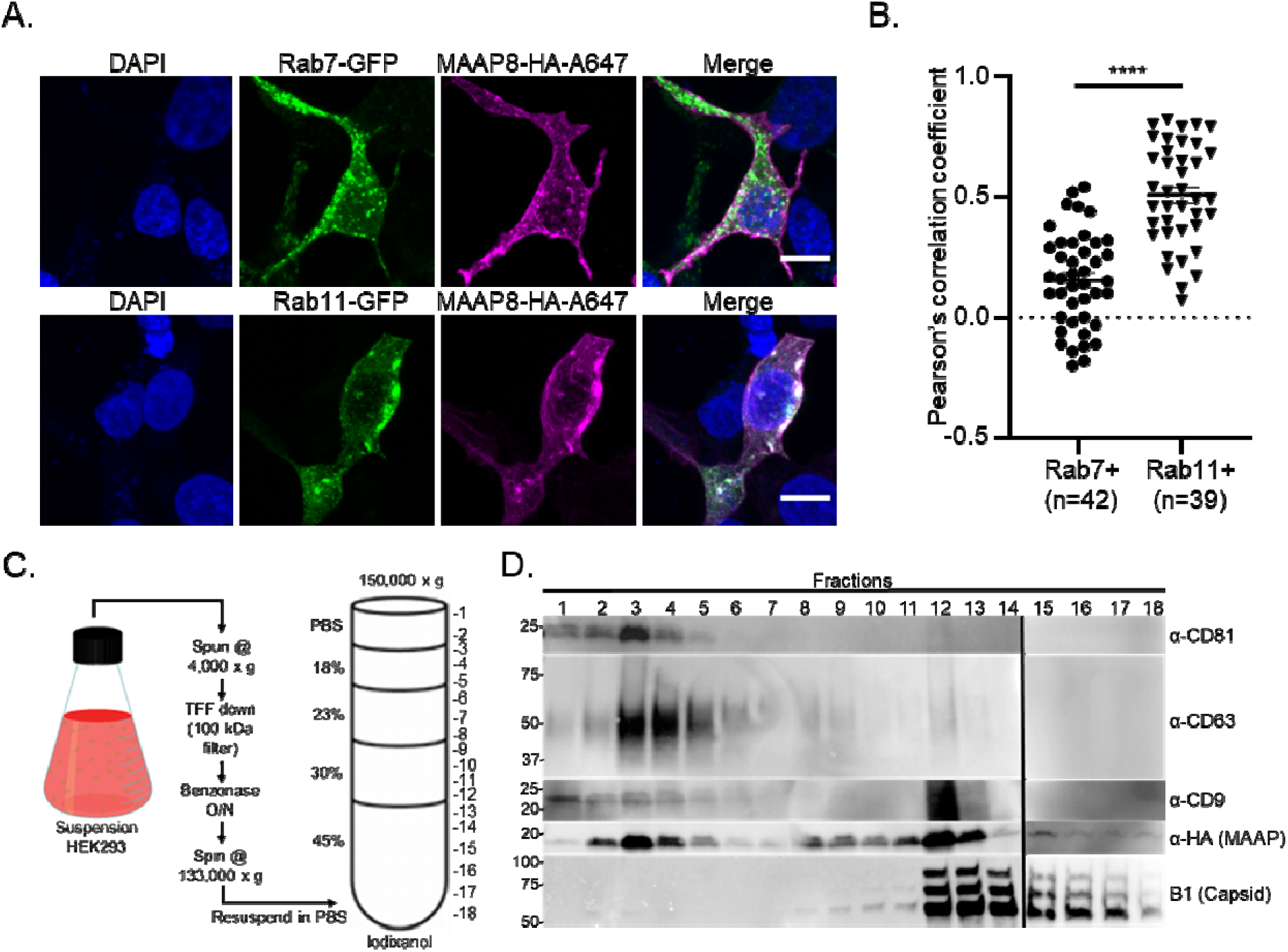
MAAP8 promotes extracellular vesicle mediated AAV cellular egress. (**A**) HEK293 cells were transfected with expression vectors encoding Rab7-GFP, Rab11-GFP, MAAP8-HA. MAAP-HA was detected by immunofluorescence with an AlexaFlour647 secondary antibody (MAAP8-HA-A647). A Z-stack of confocal optical sections at 1ψm steps was acquired. A 3-μm-thick medial stack is shown. Images are representative of three experiments. Scale bars, 10 μm. **(B)** Colocalization between MAAP8-HA and Rab7-GFP or Rab11-GFP in the whole cell as assessed by Pearson’s correlation coefficient (R) as described in Materials and Methods. Each dot represents one cell. Horizontal bars represent the mean ± SEM, Mann-Whitney rank test. \*\*\*\**p* < 0.0001, ^n.s.^*p* ≥ 0.05 **(C)** Schematic of EV isolation from HEK293 suspension culture. (D) Immunoblots of iodixanol fractions from suspension cells producing recombinant MAAP8Δ vector complemented *in trans* with MAAP8-HA. EVs, capsid and MAAP were analyzed from the media of HEK293 producing cells at day 3 post transfection. Capsid and MAAP proteins were analyzed by SDS-PAGE under reducing conditions while EV markers (CD81, CD63, CD9) were analyzed by SDS-PAGE under non-reducing conditions.

To further explore the biology of the MAAP-dependent AAV secretory process, we adopted a gradient centrifugation method to purify EVs from large volumes of cell culture supernatant^28,29^. Cell culture medium from suspension adapted HEK293 rAAV8/MAAPΔ producing cells (complemented in *trans* with MAAP8-HA) was processed by successive filtration and centrifugation steps to generate a crude EV pellet that was then separated on an iodixanol gradient **(Fig. 5C)**. The gradient was split into 18 different fractions and probed for EV, MAAP and capsid specific antibodies. Using SDS-PAGE and immunoblot analysis, we found that MAAP associated with multiple EV containing fractions and also strongly associated with an EV/AAV containing fraction **(Fig. 5D)**. To further understand this interaction between MAAP and extracellular vesicles, we purified EVs from the extracellular fractions of rAAV8 or rAAV8/MAAPΔ producing cells using the ExoQuick-TC kit^30,31^. In contrast to the rAAV8 control, we found negligible association of AAV particles associated with the purified EV fraction in the absence of MAAP as evidenced by western blot analysis of capsid proteins **(Supplementary Fig. 7A)**. Strikingly, we also observed a markedly lower signal for CD81^32,33^, a marker for EVs in the rAAV8/MAAPΔ samples compared to rAAV8 controls **(Supplementary Fig. 7A)**. These results further led us to explore the function of MAAP alone, unrelated to AAV biology. First, we observed that overexpression of MAAP-HA when compared to an HA tag only control yielded a significantly higher proportion of CD63+ EV fraction (**<u>Supplementary</u> Fig. 7B**). Second, when overexpressing MAAP8-GFP or a GFP only control, the former was significantly enriched in the purified EV fraction, directly corroborating the role of MAAP in promoting the secretion and loading of cargo into EVs (**<u>Supplementary</u> Fig. 7C**). Lastly, ultrastructural characterization of purified EV fractions using negative stain transmission electron microscopy (TEM) under different conditions (rAAV8, rAAV8/MAAPΔ, rAAV8/MAAPΔ + MAAP transcomplementation) revealed striking differences in the morphology, homogeneity and abundance of EVs **(Supplementary Fig. 8)**. In particular, MAAP overexpression appears to promote a notable increase in spherical vesicles with a diameter ranging from ~20-50 nm. Interestingly, these vesicles appear to be relatively uniform and homogenous in their composition indicating MAAP may promote secretion of a specific type of extracellular vesicle.

## Discussion

Dissecting new aspects of AAV biology is critical towards continued improvement of recombinant vectors for gene therapy applications. For instance, our lab and others previously dissected the functional attributes of the Assembly Activating Protein (AAP), which is encoded from an alternative ORF in the AAV *cap* gene^4,5,34^. These studies provided mechanistic insight into AAV capsid assembly in addition to highlighting capsid mutations that can adversely impact assembly and vector yield. Here, we provide structural and functional insights into the recently identified Membrane Associated Accesory Protein (MAAP), which is encoded from a different, alternative ORF within the AAV *cap* gene. We report that this highly conserved protein displays critical hydrophobic and cationic domains essential for membrane anchoring and promoting extracellular secretion of AAV particles. Although attenuation of MAAP expression does not affect viral yield, we demonstrate that extracellular secretion of AAV particles is drastically delayed with increased cellular retention of AAV particles. Our findings have immediate implications for recombinant AAV vector production, which is currently undergoing a revolutionary manufacturing scale-up phase^7,35^. Specifically, we demonstrate that MAAP transcomplementation can significantly alter the kinetics of AAV extracellular secretion potentially facilitating downstream purification only from cell culture media. Further, we note that mutation of the *cap* gene in engineered AAV capsids within the overlapping MAAP reading frame could impact vector yield and timing of harvest from cell lysate versus media fractions. Interestingly, we observed a significant titer increase on Day 3 with MAAP1Δ, MAAP8Δ and MAAP9Δ vectors. One possible explanation for the increased titer is that in cells transfected with MAAPΔ plasmids, empty rAAV capsids are not as readily secreted into the media and are retained in the nucleus, thereby increasing their availability for REP mediated genome packaging. In addition, we also observed that MAAP contributes to rapid secretion of what appears to be a homogenous EV fraction independent of AAV components. While this aspect needs to be explored further, it is exciting to note that MAAP might provide an orthogonal solution to loading therapeutic cargo into EVs.

From a structural perspective, MAAP is rich in alpha helical domains interspersed by T/S rich linker regions and basic residues. The most notable attribute of MAAP is a cationic amphipathic C-terminal domain, which is critical for membrane anchoring and extracellular secretion. Recent studies have revealed that PTGFRN and BASP1 can be utilized as scaffolds for loading various molecules onto the surface or into the lumen of EVs^29^. BASP1 is an inner membrane leaflet localized protein that utilizes a polybasic effector domain (PED) to bind to the EV membrane. Interestingly, the PED along with N-terminal myristoylation of BASP1 is essential for proper loading of carrier proteins to the inner membrane of EVs. Similarly, the matrix (MA) membrane proximal domain of HIV-1 Pr55^GAG^ contains a highly basic region that is important for efficient membrane binding and proper targeting of MA to the plasma membrane^36^. MAAP also contains multiple basic regions that are essential for proper molecular function; however whether MAAP undergoes post-translational lipid modification remains to be determined. Nevertheless, our data suggests that similar to BASP1, MAAP can potentially be fused to protein cargo for loading into EVs.

Furthermore, our work has shed light on the interaction between MAAP, AAV and EVs. Using established MISEV criteria^37^, our work revealed that secreted AAV particles associate with multiple EV markers (CD81, CD63 and CD9), although a fraction of AAV particles appears to be free of any vesicular association. Notably, the EV fractions showed strong MAAP association further implicating this viral protein in vesicle mediated AAV cellular egress. Whether the free particles were previously associated with EVs that lysed or were secreted independently remains the subject of investigation. From a cell biology perspective, our data supports a model wherein MAAP likely exploits molecular interactions within the EV/exosomal pathway. Exactly how MAAP exploits this pathway and how AAV particles exploit these subcellular organelles prior to secretion warrants further investigation. It is plausible that MAAP may enable passive loading of AAV particles into EVs during secretion or alternatively, actively recruit AAV capsids into EVs through an unknown cellular bridging factor. These hypotheses are further corroborated by a consistent colocalization of AAV particles with EV markers and preliminary evidence from proximity ligation. Indeed studies exploring cellular egress of autonomous parvoviruses, such as Minute Virus of Mice (MVM) have shown viral particles to be sequestered into COPII-vesicles in the endoplasmic reticulum (ER) and transported via the Golgi compartment to the plasma membrane^38^. Remodeling of the cytoskeletal filaments through gelsolin mediated actin fiber degradation in infected cells have been shown essential for egress. Further, several biochemical factors such as sar1, sec24, rab1, the ERM family proteins, radixin and moesin have been implicated in this process of cellular exocytosis^39^. However, it should be noted that a virulence factor akin to MAAP in other parvoviruses has not been identified to date. In summary, we conclude that while dependoparvoviruses such as AAV rely on helper coinfection to trigger viral release through a lytic process, cellular egress likely accelerates and regulates viral dissemination and spread. In summary, our studies unequivocally implicate MAAP as a novel AAV egress factor.

## Methods

### Plasmid Constructs

MAAP DNA sequences from AAV serotypes 1,2,5,8 and 9 were synthesized and cloned into pcDNA3.1(+)-C-HA and pcDNA3.1(+)-C-eGFP expression vectors using HindIII and XbaI sites for MAAP 1,2,8,9 and EcoRV sites for MAAP5 (Genscript). All MAAP expression constructs were synthesized and cloned with an ATG start codon. The AAV8-Rep/Cap-VP* plasmid is a AAV2-Rep/AAV8-Cap plasmid with the start codons of VP1/2/3 and AAP mutated by site directed mutagenesis to prevent expression. The AAV8-Rep/Cap-MVP* additionally has a mutated MAAP start codon to prevent MAAP expression. The AAV8-MAAPΔ plasmid is a 2 AAV2-Rep/AAV8-Cap plasmid with the start codons of MAAP mutated by site directed mutagenesis to prevent expression. The AAV8-MAAP-3X-FLAG plasmid was generated by utilizing site-directed mutagenesis to incorporate a 3X-FLAG tag in frame onto the C-terminus of MAAP8 in the AAV8-Rep/Cap-VP* plasmid. AAV8-MAAP-3X-FLAG truncation mutants were generated through site directed mutagenesis. MAAP8 was cloned into the MCS-13X-Linker-BioID2-HA expression vector (Addgene #80899) using NheI and Age1 sites. All plasmid constructs were verified by DNA sequencing analysis.

### Bioinformatics analysis and structural models

The amino acid sequences of 15 AAV serotypes were retrieved from GenBank. MAAP start and stop sites were defined as previously described^18^. Protein sequences were aligned using the ClustalW multiple-alignment tool^40^ and generated using Unipro UGENE software^41^. MAAP amino acid sequences from multiple AAV isolates were aligned using ClustalW, and phylogenetic trees were generated using the MEGAv7.0.21 software package^42^. The phylogeny was produced using the neighbor-joining algorithm, and amino acid distances were calculated using a Poisson correction^43^. Statistical testing was done by bootstrapping with 1,000 replicates to test the confidence of the phylogenetic analysis and to generate the original tree^44^. The percentage of replicate trees in which associated taxa clustered together in the bootstrap test is displayed next to the branches. Secondary structural elements were predicted using the JPred tool^45^. To predict membrane-binding, amphipathic α-helices, we used Amphipaseek (parameters: high specificity/low sensitivity)^46^. MAAP structural models were generated using the TrRosetta deep learning based modeling method^19^. Secondary structural depictions of these models were visualized using the PyMOL Molecular Graphics System (Schrödinger; https://www.pymol.org/2/).

### Cellular assays, Immunoprecipitations and Western Blotting

For protein expression analysis, HEK293 cells seeded overnight in 6-well plates at a density of 3 × 10^5^ cells per plate were transfected with a total of 2 μg DNA as indicated. HEK293 cell pellets overexpressing MAAP-HA, MAAP-GFP or expressing MAAP8-3X-FLAG were recovered 72 h post-transfection. Pellets were lysed in RIPA buffer with 1x Halt Protease Inhibitor (ThermoFisher) for 45 minutes at 4°C. Lysates were spun at max speed for 10 minutes at 4°C to remove cellular debris. 1X LDS sample buffer with 10mM DTT were added to cleared lysates and boiled for 2 min. Samples of cleared lysate were ran on Mini-Protean TGX 4-15% gels (Biorad), transferred onto PVDF with the Trans-Blot Turbo system (BioRad) and blocked in 5% milk/1x TBST. Blots were probed with mouse monoclonal anti-GFP antibody (1:1000 dilution, SC9996; Santa Cruz Biotechnology), rabbit polyclonal anti-HA SG77 antibody (1:1000 dilution, 71-5500; ThermoFisher Scientific), mouse monoclonal B1 hybridoma supernatant (1:50, 03-65158; ARP), mouse monoclonal anti-FLAG M2 antibody (1:1000 dilution, F180450UG; Sigma), mouse monoclonal anti-beta actin (1:1000 dilution, 8226; Abcam) as the primary antibody. Following three 1x TBST washes, samples were incubated with secondary antibodies conjugated to HRP at 1:20,000 in 5% milk/1x TBST for 1 hour (goat anti-mouse-HRP, 32430; ThermoFisher Scientific and goat anti-rabbit-HRP, 111-035-003; Jackson ImmunoResearch). Blots were developed using SuperSignal West Femto substrate (ThermoFisherScientific/Life Technologies) according to manufacturer instructions. For immunoprecipitation studies, HEK293 cells were transfected with pXR9 and MAAP9-pcDNA3.1(+)-C-HA for 72 hrs, then washed with 1X PBS and harvested in NP-40 with 1x Halt Protease Inhibitor (ThermoFisher) for 1 hr at 4°C. Lysates were spun at max speed for 20 minutes at 4°C to remove cellular debris. 10μL (2.5μg) of anti-HA SG77 antibody were added to 500 μL cleared lysate and incubated at 4°C for 3 hrs with nutation. Added 40 μL of pre-washed Protein G magnetic beads to pre-cleared lysate with antibody and carried out immunoprecipitations overnight on a nutator at 4°C. Bound protein was eluted in 10mM DTT and 1x LDS for 5 min at 95°C. Samples were then analyzed via SDS-PAGE (NuPAGE 4-12% Bis-Tris Gel) and transferred onto nitrocellulose membrane (ThermoScientific). Following blocking in 5% milk/1xTBST, samples were incubated with primary antibodies to either capsid (mouse monoclonal B1 hybridoma supernatant, 1:250, 65158; Progen), actin (mouse monoclonal anti-beta actin, 1:1000 dilution, 8226; Abcam), CD81 (mouse monoclonal anti CD81 M38, 1:1000, 10630D), CD63 (mouse monoclonal anti CD63 Ts63, 1:1000, 10628D), CD9 (mouse monoclonal anti CD9 Ts9, 1:1000, 10626D) or MAAP (mouse monoclonal anti-HA HA.C5 antibody, 1:1000 dilution, MA5-27543; ThermoFisher Scientific) overnight in 5% milk/1xTBST. Following three 1x TBST washes, samples were incubated with secondary anti-mouse antibody conjugated to HRP (goat anti-mouse-HRP, 32430; ThermoFisher Scientific) at 1:20,000 in 5% milk/1xTBST for 1 hour. The signal was the visualized via SuperSignal West Femto Maximum Sensitivity substrate (ThermoScientific) according to manufacturer instructions.

### BioID2 expression and pulldown

For protein expression analysis, HEK293 cells seeded overnight in 6-well plates at a density of 3 × 10^5^ cells per plate were transfected with a total of 2 μg of 13X-BioID2-HA or MAAP8-13X-BioID2-HA DNA. Cells were then supplemented with 50 μM biotin 24 hrs post transfection and allowed to label for 20 hrs. Cell pellets were recovered 20 hrs biotin supplementation and were lysed in RIPA buffer with 1x Halt Protease Inhibitor (ThermoFisher) for 45 minutes at 4°C. Lysates were spun at max speed for 10 minutes at 4°C to remove cellular debris. 1X LDS sample buffer with 10mM DTT were added to cleared lysates and boiled for 2 min. Samples of cleared lysate were ran on Mini-Protean TGX 4-15% gels (Biorad), transferred onto PVDF with the Trans-Blot Turbo system (BioRad) and blocked in 5% milk/1x TBST. Blots were probed with rabbit polyclonal anti-HA SG77 antibody (1:1000 dilution, 71-5500; ThermoFisher Scientific), mouse monoclonal anti-beta actin (1:1000 dilution, 8226; Abcam), and goat polyclonal anti-biotin (1:1000 dilution, 31852; ThermoFisher) as the primary antibody. Following three 1x TBST washes, samples were incubated with secondary antibodies conjugated to HRP at 1:20,000 in 5% milk/1x TBST for 1 hour (goat anti-mouse-HRP, 32430; ThermoFisher Scientific, goat anti-rabbit-HRP, 111-035-003; Jackson ImmunoResearch, mouse anti-goat-HRP, 31400, ThermoFisher).

Streptavidin pulldowns were performed as previously described^47^ with the following changes. HEK293 cells for each condition were grown on 2 x 15 cm^2^ plates until ~70% confluent and were transfected with 34 μg of plasmid DNA per plate. Each plate was transfected with 13X-BioID2 or MAAP8-13X-BioID2 along with pXX680, pTR-CBA-Luciferase, and AAV8-MAAPΔ. Media was supplemented with 50 μM biotin 48 hours post transfection and cells were harvested 20 hours post biotin supplementation. Biotinylated proteins were pulled down with Pierce High Capacity Streptavidin Agarose (20357; ThermoFisher) and were separated by SDS-PAGE and visualized using a SilverXpress silver stain kit (LC6100; ThermoFisher) according to the manufacturer’s instructions. Biotinylated proteins were also visualized by immunoblot and probed with rabbit polyclonal anti-HA SG77 antibody (1:1000 dilution, 71-5500; ThermoFisher Scientific), mouse monoclonal anti-beta actin (1:1000 dilution, 8226; Abcam), goat polyclonal anti-biotin (1:1000 dilution, 31852; ThermoFisher) and mouse monoclonal B1 hybridoma supernatant (1:50, 03-65158; ARP) as the primary antibody.

### Recombinant and wild type AAV production, purification, and quantification

HEK293 (human embryonic kidney cells obtained from the University of North Carolina Vector Core) were maintained in Dulbecco’s Modified Eagle’s Medium (DMEM) supplemented with 10% fetal bovine serum (FBS), 100U/ml penicillin, 100ug/ml streptomycin. Cells were maintained in 5% CO_2_ at 37°C. Recombinant AAV vectors were produced by transfecting HEK293 cells at ~75% confluence with polyethylenimine (PEI Max; Polysciences, 24765) using a triple plasmid transfection protocol with the AAV Rep-Cap plasmid, Adenoviral helper plasmid (pXX680), and single-stranded genomes encoding firefly luciferase driven by the chicken beta-actin promoter (ssCBA-Luc) or self-complementary green fluorescence protein driven by a hybrid chicken beta-actin promoter (scCBh-GFP), flanked by AAV2 inverted terminal repeat (ITR) sequences. Viral vectors were harvested from media and purified via iodixanol density gradient ultracentrifugation followed by phosphate buffered saline (PBS) buffer exchange. Titers of purified virus preparations were determined by quantitative PCR using a Roche Lightcycler 480 (Roche Applied Sciences, Pleasanton, CA) with primers amplifying the AAV2 ITR regions (forward, 5’-AACATGCTACGCAGAGAGGGAGTGG-3’; reverse, 5’-CATGAGACAAGGAACCCCTAGTGATGGAG-3’) (IDT Technologies, Ames IA).

### Quantitative PCR analysis of AAV vector yield

HEK293 cells in six-well plates were transfected using PEI at ~75% confluence with Adenovirus helper plasmid (1μg), WT or MAAPΔ AAV-Rep/Cap plasmid (1μg), ITR-transgene plasmid (500ng), and AAV8-Rep/Cap-VP* or -MVP* (500ng). The AAV8-Rep/Cap-VP* plasmid is a AAV2-Rep/AAV8-Cap plasmid with the start codons of VP1/2/3 and AAP mutated to prevent expression. The AAV8-Rep/Cap-MVP* additionally has a mutated MAAP start codon. For 3-day experiments, media and cells were collected three days post-transfection. Cells were lysed by vortexing in a mild lysis buffer (10mM Tris-HCl, 10mM MgCl_2_, 2mM CaCl_2_, 0.5% Triton X-100 supplemented with DNAse, RNAse, and Halt Protease Inhibitor Cocktail) and incubated at 37°C for 1hr. Lysates were cleared by centrifuging at 21,000 rcf for two minutes. NaCl to 300mM was added to AAV2 lysates prior to centrifugation to prevent virus binding of the cell debris. Collected media and cleared lysates were assayed with qPCR for DNAse-resistant viral genomes as described above. For Day 3 and 5 experiments, media was collected and replaced on the first indicated day, and cells and media were harvested as described on the last day. For WT-like AAV8 transfections, cells were transfected with Adenovirus helper plasmid (1.9μg) and WT or MAAPΔ ITR-AAV2-Rep/AAV8-Cap-ITR plasmid (0.9μg), with media and cells collected on days 3 and 5 post-transfection as described above.

### Confocal fluorescence microscopy

HEK293 cells were seeded on slide covers in 24-well plates at a density of 5e4 cells/well and allowed to adhere overnight. Cells were then co-transfected with Rab7-GFP, Rab11-GFP, MAAP8-pcDNA3.1(+)-C-HA and MAAP9-pcDNA3.1(+)-C-HA, and were incubated for 48 hrs at 37°C and 5% CO_2_. Cells were then fixed with 4% paraformaldehyde for 30 minutes and permeabilized with 0.1% Triton X-100 for 30 minutes. Following 1 hr of blocking with 5% Normal Goat Serum, cells were stained with rabbit polyclonal anti-HA SG77 antibody (1:100 dilution, 71-5500; ThermoFisher Scientific) primary for one hour, washed 3x with PBS, and then stained with fluorescent goat anti-rabbit Alexa Fluor 647 (1:400 dilution, ab150079; Abcam) secondary antibody and washed 1x with PBS. Cells were subject to 5 minutes staining with DAPI, and then mounted in Prolong Diamond (Invitrogen) and imaged using a Zeiss LSM 880 Airyscan confocal microscope. Co-localization analysis was performed by cropping the whole compartment (Rab11 or FIP3), or the whole cell, using Zeiss ZEN software with the Colocalization function. Threshold was automatically determined using the Costes method autothreshold determination. Pearson’s correlation coefficient was calculated for the analysis. Statistical analyses were carried out by the nonparametrical Mann–Whitney *U* test using Prism software (GraphPad).

### Extracellular vesicle Isolation

For Figures 5C-D and Supplementary Figure 7D-H, a previously established EV isolation protocol was utilized^29^. Suspension adapted HEK293 (human embryonic kidney cells obtained from the University of North Carolina Vector Core) cells were grown in Dulbecco’s Modified Eagle’s Medium (DMEM) supplemented with 10% fetal bovine serum (FBS), 100U/ml penicillin, 100ug/ml streptomycin and inoculated at a density of 5E+5 viable cells per milliliter. Cells were maintained in 5% CO_2_ at 37°C and grown overnight to a density of 1E+6 cells per milliliter for transfection. Cells were then grown for 3 days with viability measured daily. Cell density and viability were measured on a Countess II Automated Cell Counter (Invitrogen). Upon culture termination, cells were removed by centrifugation at 4,000 x g for 30 minutes at room temperature. Harvests were then concentrated using tangential flow filtration using a Minikros (100 kD) modified polyethersulfone (PES) hollow fiber filter (Repligen) on a KROSFLO KMPI system. Concentrated medium was then supplemented with 1mM MgCl_2_ and benzonase (20 U/mL, Sigma Aldrich) to digest extravesicular nucleic acids. Nuclease treated media was next transferred to 38-mL Ultra-Clear centrifuge tubes (Beckman Coulter) and centrifuged for 60 min at 133,000 x g at 4°C in a Sure Spin 630 36 mL rotor (Thermo Scientific). The media was discarded and the crude pellet was resuspended in a minimal volume of PBS. The resuspended pellet was then mixed with a 60% iodixanol solution (Optiprep, Sigma) and layered onto the bottom of a 38-mL Ultra-Clear centrifuge tube (Beckman Coulter). Lower-density solutions were prepared by diluting homogenization buffer (250 mM sucrose, 10 mM Tris-HCl, 1 mM EDTA, pH 7.4) to yield final iodixanol concentrations (vol/vol) of 45%, 30%, 23%, and 18%. Successive layers of 30% (9 mL), 23% (6 mL) and 18% (6 mL) iodixanol solutions were carefully pipetted on top. Resuspended pellet in PBS was added on top of the gradient. The gradient was ultracentrifuged in a Sure Spin 630 36 mL rotor (Thermo Scientific) for 16 h at 150,000 x g and 4°C to separate EVs from other cell culture supernatant contaminants. The gradient was then fractionated in 2 mL fractions and analyzed by biochemical analysis. For Supplementary Figure 7A-C and Supplementary Figure 8, EVs were isolated from tissue culture media using a commercial kit containing a polyethylene glycol (PEG) solution (ExoQuick-TC ULTRA kit, EQULTRA-20TC-1; System Biosciences) according to the manufacturer’s instructions.

### Transmission electron microscopy

Isolated vesicular fractions or AAV viral particles (1×10^10^ vg) in 1x PBS samples were adsorbed onto 400 mesh, carbon coated grids (Electron Microscopy Sciences) for 2 min and briefly stained with 1% uranyl acetate (Electron Microscopy Sciences) diluted in 50% ethanol. After drying, grids were imaged with a Philips CM12 electron microscope operated at 80kV. Images were collected on an AMT camera.

### Luciferase expression assays

A total of 1×10^4^ HEK293 cells in 50μL DMEM + 10% FBS + penicillin streptomycin was then added to each well, and the plates were incubated in 5% CO2 at 37 °C for 24h. Cells were then transduced with AAV8-ssCBA-Luc vectors at a dose of 10,000 and 50,000 vg/cell. 48 hours post transduction, cells were then harvested and lysed with 25μL of 1×passive lysis buffer (Promega) for 30 min at room temperature. Luciferase activity was measured on a Victor 3 multi-label plate reader (PerkinElmer) immediately after the addition of 25μL of luciferin (Promega).

### Statistical Analysis

Where appropriate, data are represented as mean or mean ± standard deviation. For data sets with two groups (Figure 2B and 2E; Supplemental Figure 1A), comparisons were made between all groups and significance was determined using a two-way ANOVA followed by a Sidak’s post-test. For data sets with at least three groups (Figure 3), comparisons were made between all groups and significance was determined by two-way ANOVA, with Tukey’s post-test. For analysis of confocal microscopy data (Figure 4A; Supplemental Figure 4A) significance was determined by a Mann-Whitney rank test. \**p*<0.05, \*\**p*<0.01, \*\*\**p*<0.001, \*\*\*\**p*<0.0001.

## Acknowledgements

This study was funded by NIH grants awarded to A.A. (R01HL089221, UG3AR07336, R01GM127708; R01NS099371). We would like to acknowledge Ricardo Vancini and the Duke Research Electron Microscopy Service for their help in the preparation and imaging of TEM samples. We would also like to acknowledge Dana Elmore for her assistance with data analysis.

## Author Contributions

ZE, PH, and AA designed all experiments and interpreted the data. ZE, PH and DO carried out all molecular biology, virus production and imaging studies. ZE, PH, HV and AA wrote the manuscript.

## Competing Interests

AA and ZE have filed patent applications on the subject matter of this manuscript. AA is a founder at StrideBio, TorqueBio and WardenBio.

---

**Supplementary Figure 1.**
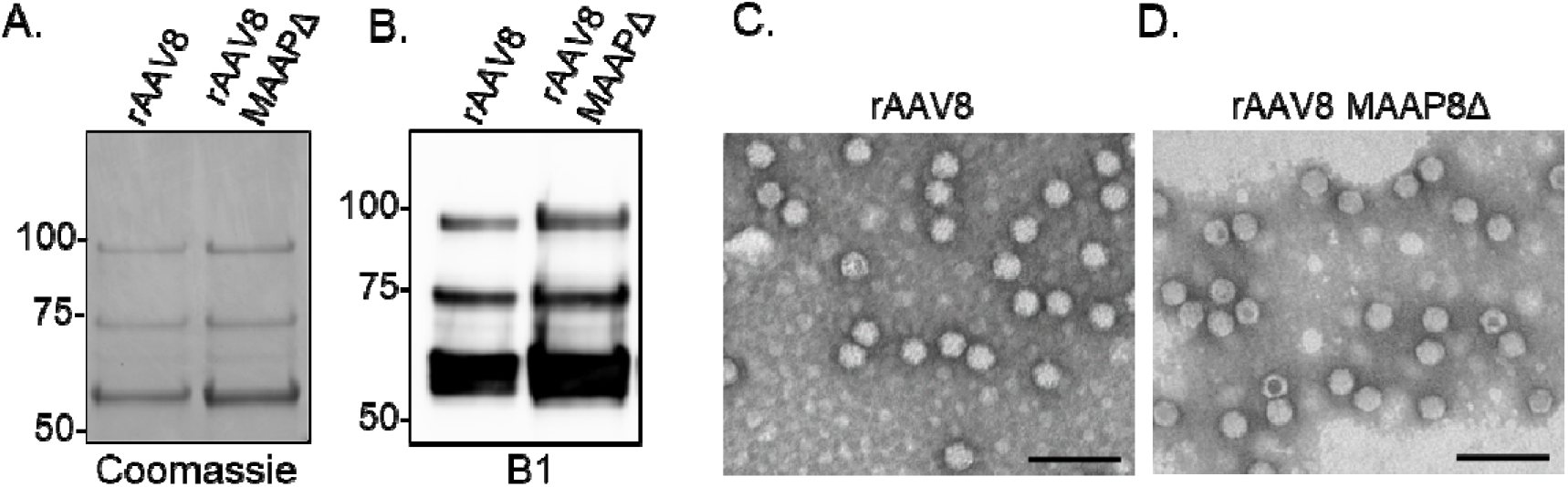
MAAP ablation does not affect AAV capsid protein composition and morphology. **(A)** Recombinant AAV8 and AAV8 MAAPΔ virus was purified from the media of HEK293 producing cells. AAV8 and AAV8 MAAPΔ viral capsids were analyzed by SDS-PAGE under reducing conditions and stained with coomassie **(A)** or probed with a capsid (B1) specific antibody **(B)**. TEM images of rAAV8 **(C)** and rAAV8 MAAPΔ **(D)** viral capsids.

**Supplementary Figure 2.**
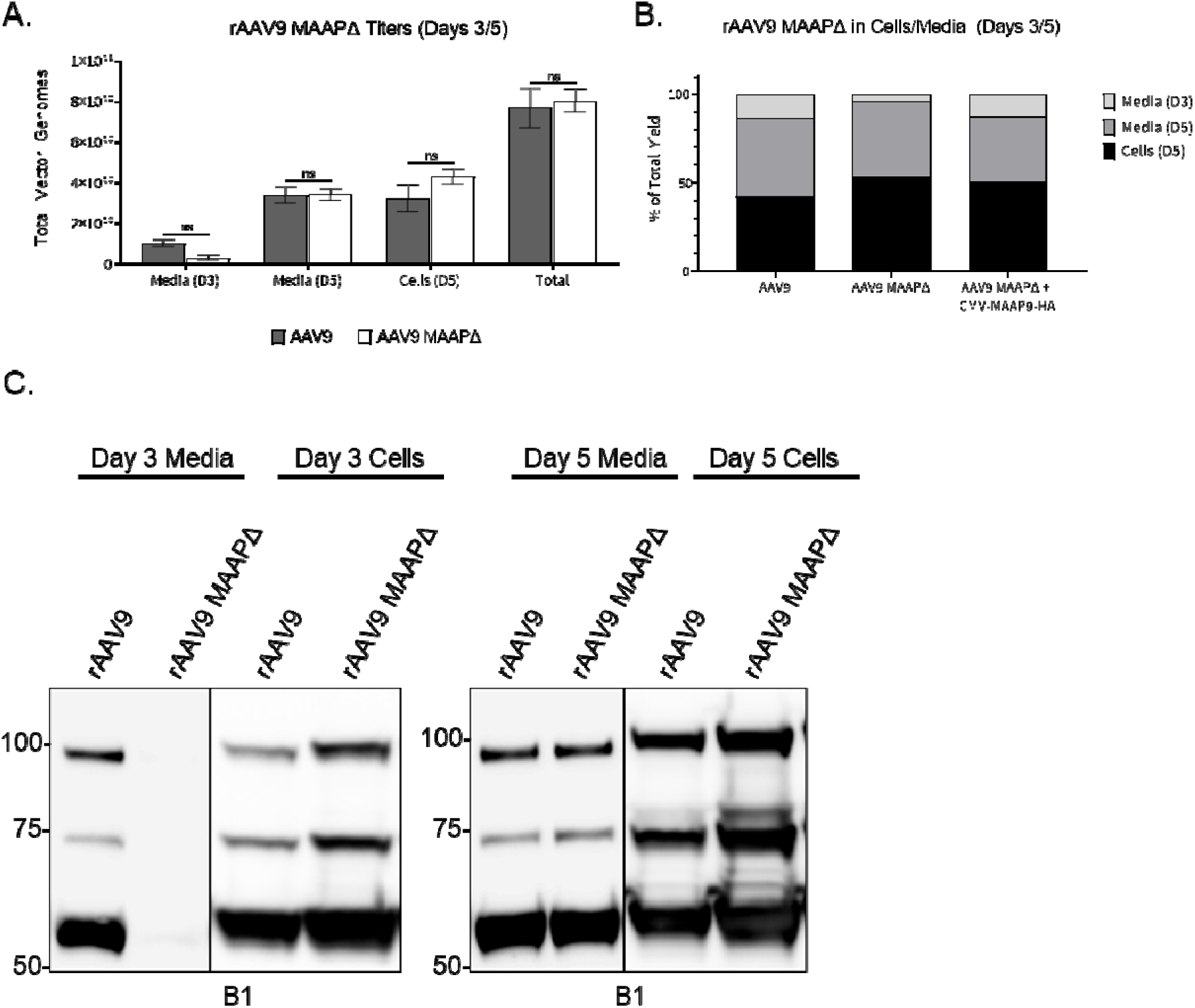
Ablation of MAAP expression differentially impacts recombinant AAV9 secretion. **(A)** AAV9 ssCBA-Luc vectors produced with WT Cap or MAAPΔ Cap. Total vector genomes collected from the cells and media and the proportion of virus found in each media harvest or associated with the cells **(B)** are shown. **(C)** Recombinant AAV9 and AAV9 MAAPΔ viruses were analyzed from the media and pellet of HEK293 producing cells at days 3 and 5 post transfection. Capsid proteins were analyzed by SDS-PAGE under reducing conditions and probed with a capsid (B1) specific antibody.

**Supplementary Figure 3.**
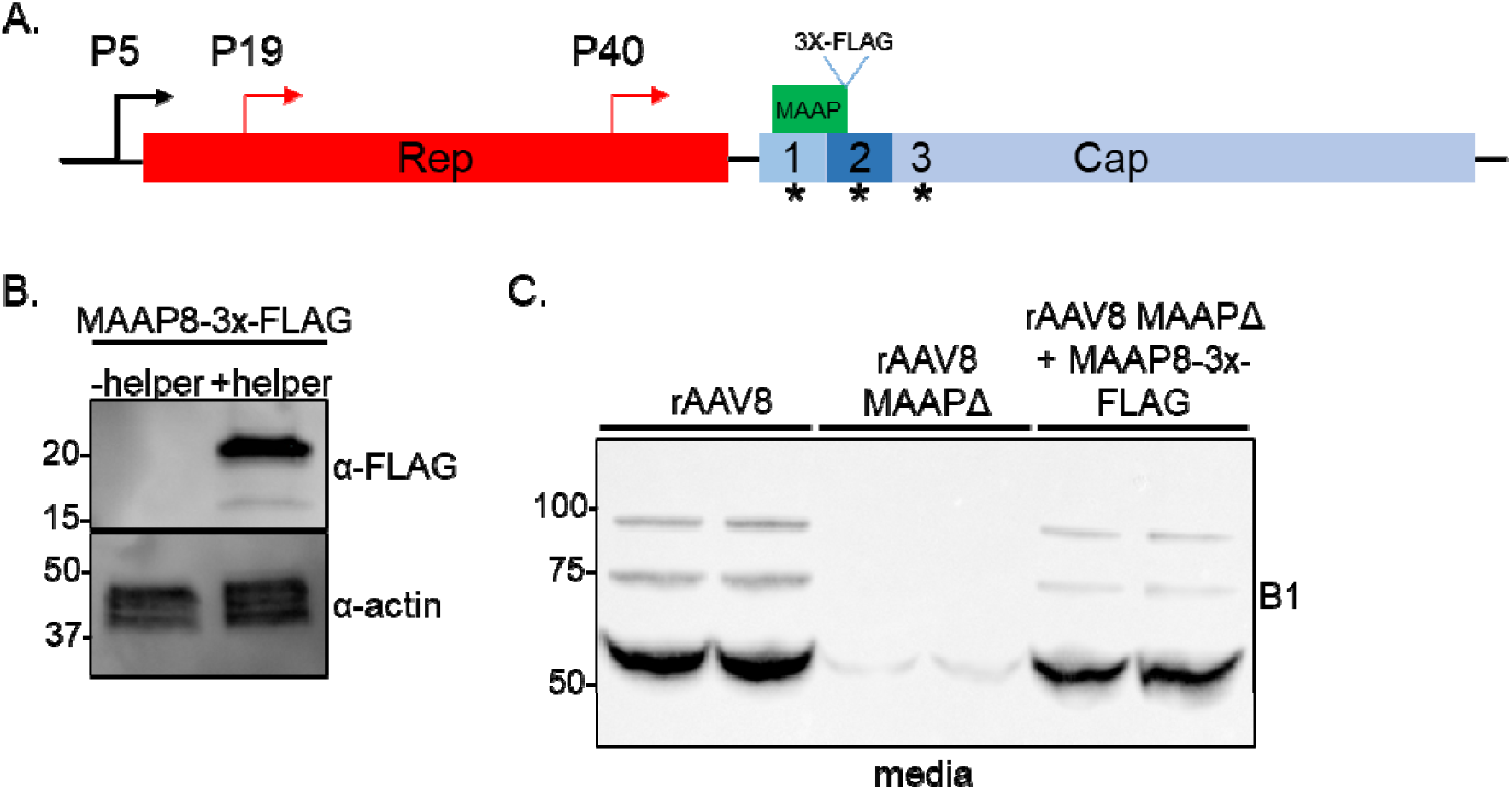
MAAP8-3X-FLAG rescues MAAP8Δ function. **(A)** Schematic of recombinant AAV8 VP/AAP-null MAAP8-3X-FLAG (MAAP8-3X-FLAG) plasmid used to replicate endogenous levels of MAAP expression. **(B)** HEK293 cells were transfected with MAAP8-3X-FLAG along with pXX680 (Adenoviral helper) plasmids and harvested 72 hours post transfection. Whole cell lysate was analyzed by SDS-PAGE under reducing conditions and probed with FLAG (α-FLAG) and actin (α-actin) specific antibodies. **(C)** Recombinant AAV8 and AAV8 MAAPΔ viruses complemented with MAAP8-3X-FLAG were analyzed from the media of HEK293 producing cells at day 3 post transfection. Capsid proteins were analyzed by SDS-PAGE under reducing conditions and probed with a capsid (B1) specific antibody.

**Supplementary Figure 4.**
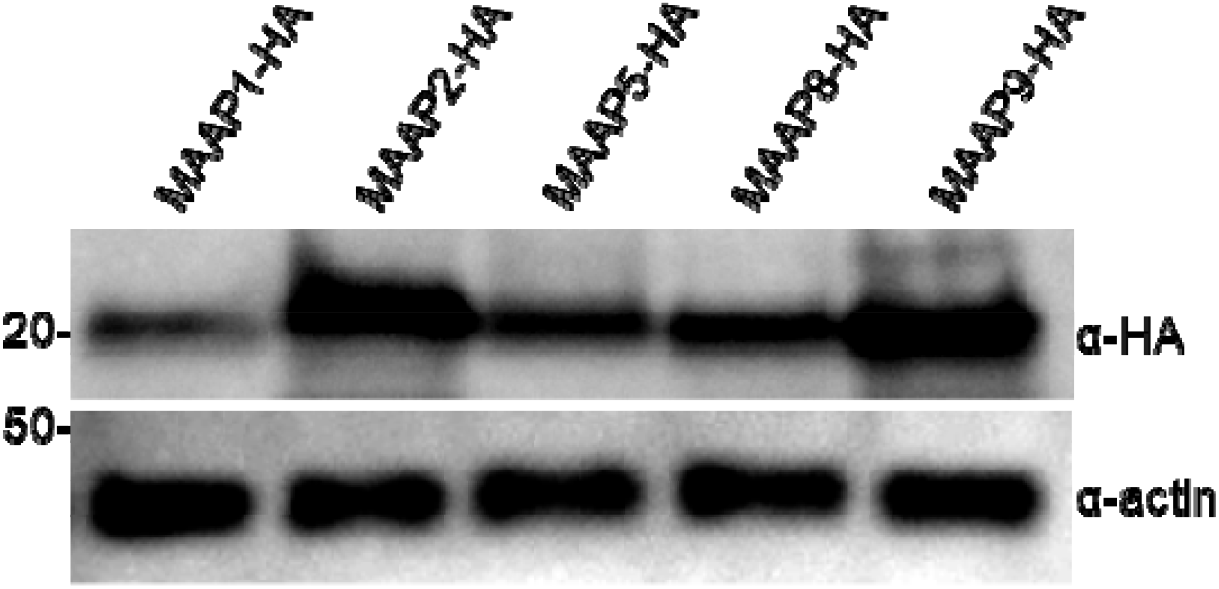
Expression of MAAP-HA fusions. Anti-HA immunoblot of wholecell extracts prepared from HEK293 cells expressing indicated HA tagged constructs. Anti-actin immunoblot served as loading control.

**Supplementary Figure 5.**
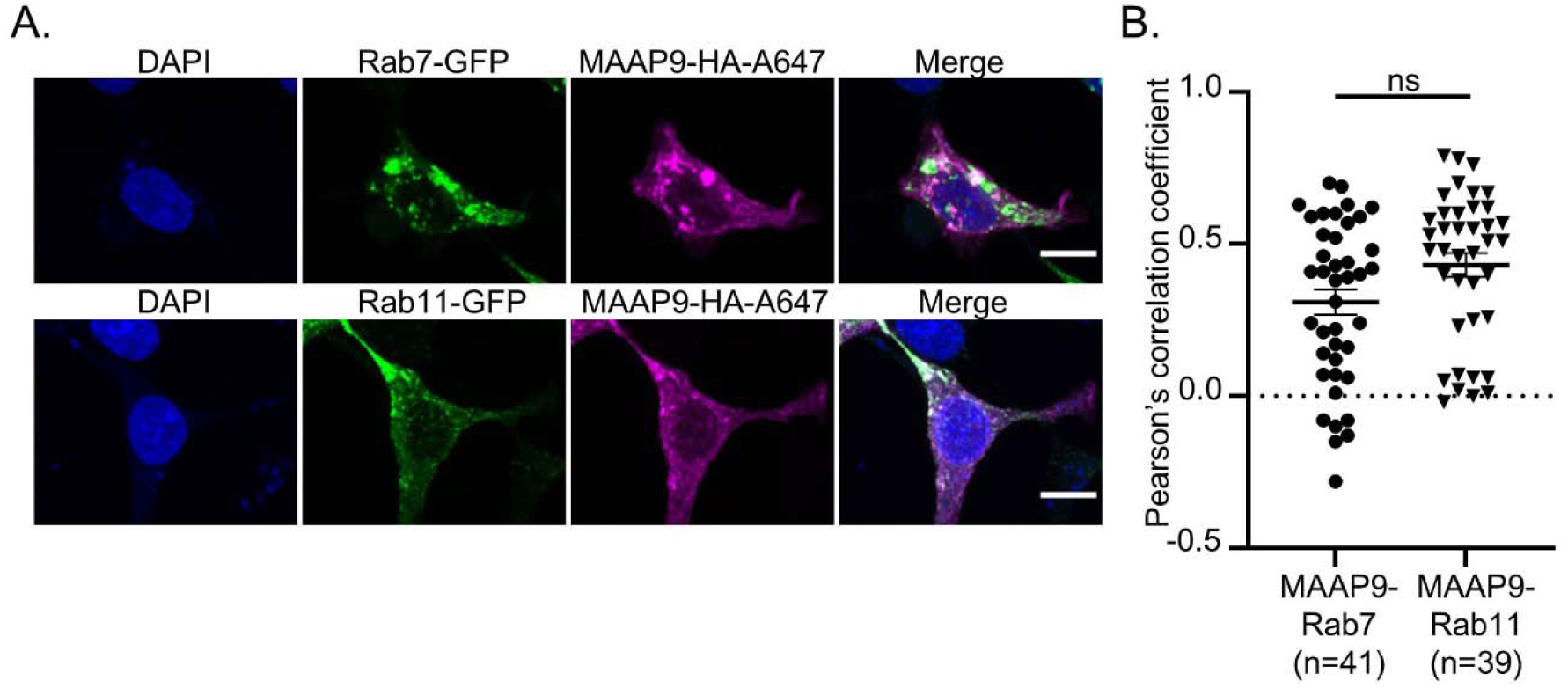
MAAP9 demonstrates similar colocalization with endocytic and exocytic vesicular markers. **(A)** HEK293 cells were transfected with expression vectors encoding Rab7-GFP, Rab11-GFP, MAAP9-HA. MAAP-HA was detected by immunofluorescence with an AlexaFlour647 secondary antibody (MAAP9-HA-A647). A Z-stack of confocal optical sections at 1-μm steps was acquired. A 3-μm-thick medial stack is shown. Images are representative of three experiments. Scale bars, 10 μm. (**B**) Colocalization between MAAP9-HA and Rab7-GFP or Rab11-GFP in the whole cell as assessed by Pearson’s correlation coefficient (R) as described in Materials and Methods. Each dot represents one cell. Each dot represents one cell. Horizontal bars represent the mean ± SEM, Mann-Whitney rank test.^n.s.^ *p* ≥ 0.05

**Supplementary Figure 6.**
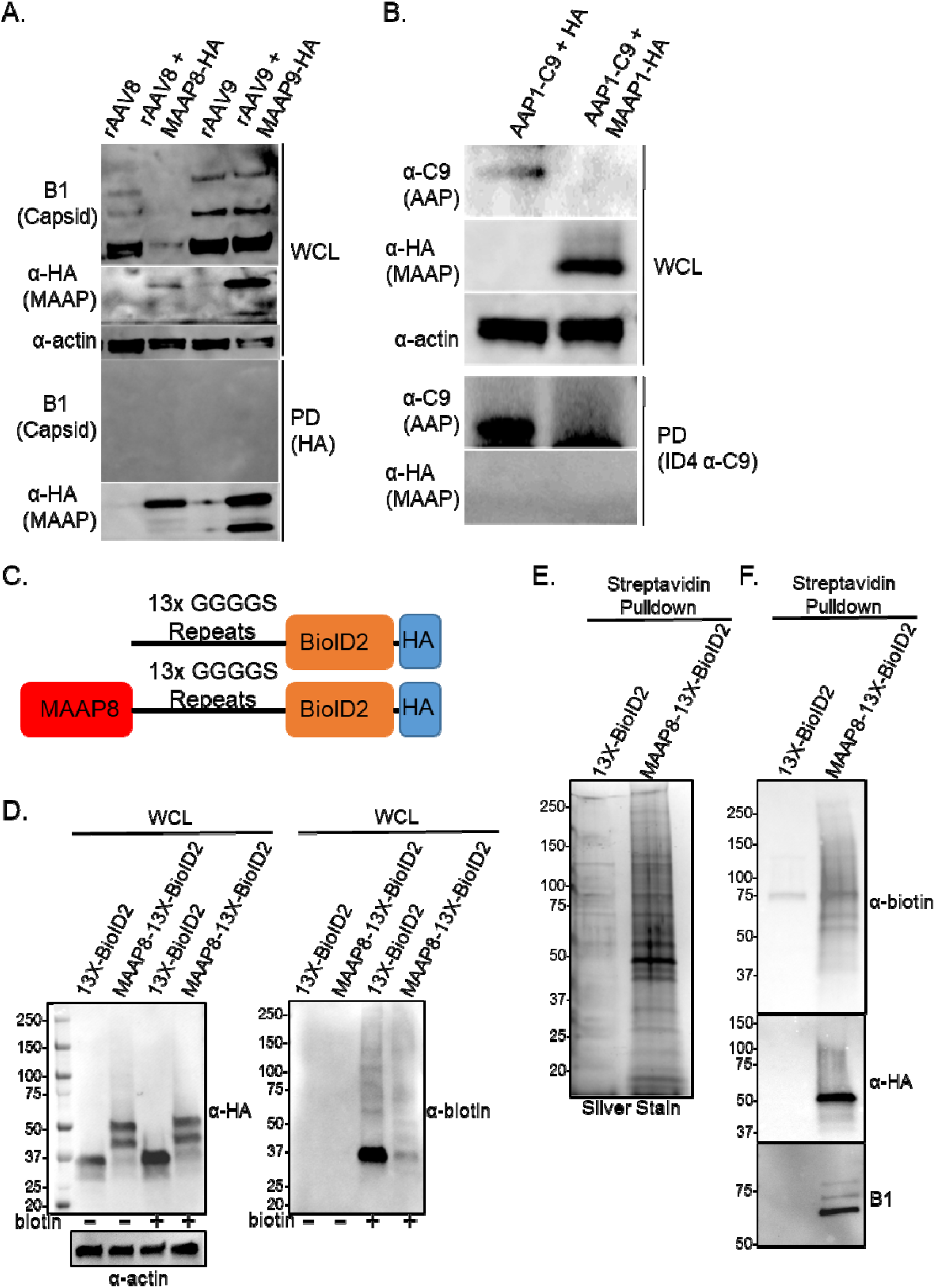
MAAP proximally interacts with the AAV capsid. **(A)** Immunoprecipitation (IP) of MAAP8/9-HA with rAAV8 and rAAV9 capsids and immunoblotting of input whole cell lysate (WCL) and pull down (PD) material for actin, capsid (B1), and MAAP8/9 (HA). **(B)** IP of AAP1-C9 and MAAP1-HA and immunoblotting of input whole cell lysate (WCL) and pull down (PD) material for actin, capsid (B1), MAAP8/9 (HA) and AAP (ID4). **(C)** Schematic of MAAP8-13X-BioID2-HA fusions. **(D)** HEK293 cells were transfected with expression vectors encoding 13X-BioID2 and MAAP8-13X-BioID2. Media was supplemented with 50 μM biotin 24 hours post transfection and cells were harvested 24 hours post biotin supplementation. Whole cell lysate (WCL) was analyzed by SDS-PAGE under reducing conditions and probed with HA (α-HA), biotin (α-biotin) and actin (α-actin) specific antibodies. **(E)** HEK293 cells were transfected with plasmids encoding either 13X-BioID2 or MAAP8-13X-BioID2 along with pXX680, pTR-CBA-Luciferase, and AAV8-MAAPΔ. Media was supplemented with 50 μM biotin 48 hours post transfection and cells were harvested 20 hours post biotin supplementation. Biotinylated proteins pulled down on streptavidin resin were separated by SDS-PAGE and visualized by silver stain or **(F)** probed with biotin (α-biotin), (α-HA), and capsid (B1) specific antibodies.

**Supplementary Figure 7.**
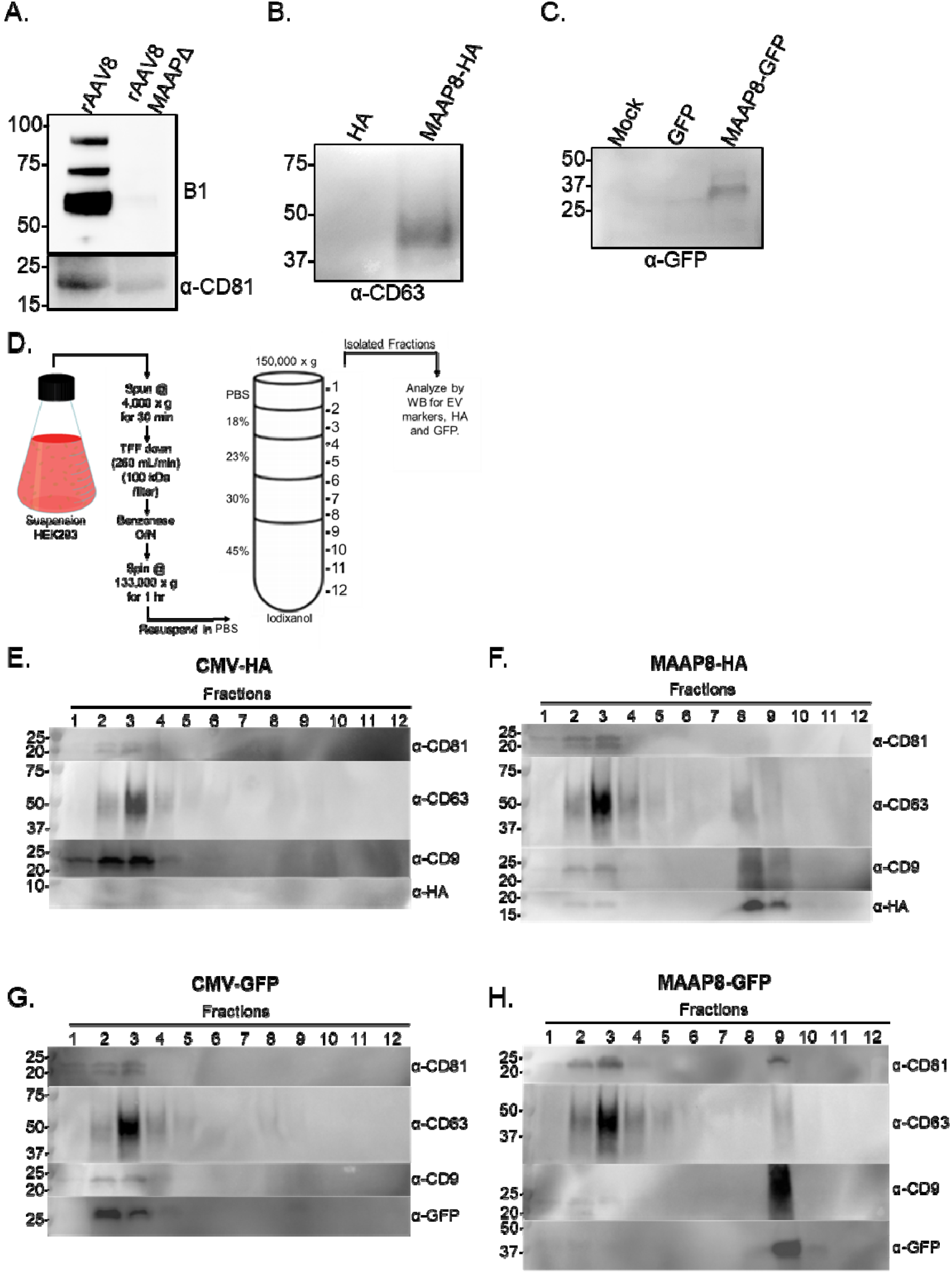
MAAP promotes secretion of extracellular vesicles. **(A)** EVs were isolated from media of AAV8 and AAV8 MAAPΔ producing HEK293 cells using the ExoQuick-TC Ultra based method. Isolated EVs were analyzed by SDS-PAGE and probed with capsid (B1) and EV (α-CD81) specific antibodies. **(B)** HEK293 cells were transfected with expression vectors encoding HA and MAAP8-HA with EVs being isolated from media 72 hours post transfection. Isolated EVs were analyzed by SDS-PAGE under non-reducing conditions and probed with EV (α-CD63) specific antibody. **(C)** HEK293 cells were transfected with expression vectors encoding GFP and MAAP8-GFP with EVs being isolated from media 72 hours post transfection. Isolated EVs were analyzed by SDS-PAGE under reducing conditions and probed with a GFP (α-GFP) specific antibody.) **(D)** Schematic of EV isolation from HEK293 suspension culture. Immunoblots of iodixanol fractions from suspension cells expressing **(E)** CMV-HA, **(F)** MAAP8-HA, **(G)** CMV-GFP and **(H)** MAAP8-GFP. EVs and MAAP were analyzed from the media of HEK293 cells at day 3 post transfection. HA and GFP proteins were analyzed by SDS-PAGE under reducing conditions while EV markers (CD81, CD63, CD9) were analyzed by SDS-PAGE under non-reducing conditions.

**Supplementary Figure 8.**
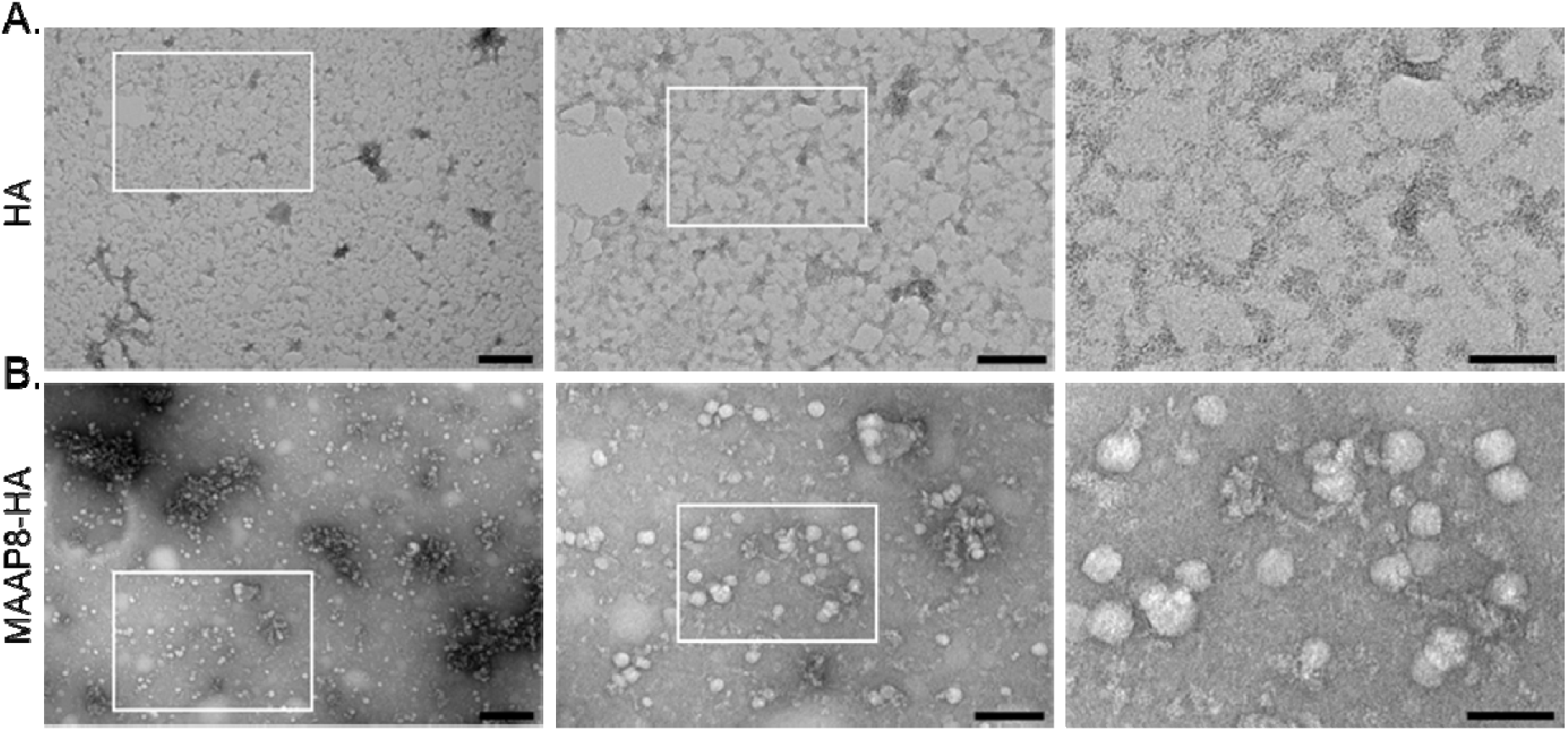
MAAP promotes EV production. **(A)** HEK293 cells producing recombinant AAV8 MAAPΔ were transfected with expression vectors encoding HA and **(B)** MAAP8-HA. EVs were isolated from media of AAV producing HEK293 cells 72 hours post transfection using an ExoQuick-TC Ultra kit and analyzed by TEM. Highlighted top inset region magnified in bottom image. Left-Scale bars, 200 nm. Middle-Scale bars, 100 nm. Right-Scale bars, 50 nm.

